# 1.4 min Plasma Proteome Profiling via Nanoparticle Protein Corona and Direct Infusion Mass Spectrometry

**DOI:** 10.1101/2024.02.06.579213

**Authors:** Yuming Jiang, Jesse G. Meyer

## Abstract

Non-invasive detection of protein biomarkers in plasma is crucial for clinical purposes. Liquid chromatography mass spectrometry (LC-MS) is the gold standard technique for plasma proteome analysis, but despite recent advances, it remains limited by throughput, cost, and coverage. Here, we introduce a new hybrid method, which integrates direct infusion shotgun proteome analysis (DISPA) with nanoparticle (NP) protein coronas enrichment for high throughput and efficient plasma proteomic profiling. We realized over 280 protein identifications in 1.4 minutes collection time, which enables a potential throughput of approximately 1,000 samples daily. The identified proteins are involved in valuable pathways and 44 of the proteins are FDA approved biomarkers. The robustness and quantitative accuracy of this method were evaluated across multiple NPs and concentrations with a mean coefficient of variation at 17%. Moreover, different protein corona profiles were observed among various nanoparticles based on their distinct surface modifications, and all NP protein profiles exhibited deeper coverage and better quantification than neat plasma. Our streamlined workflow merges coverage and throughput with precise quantification, leveraging both DISPA and NP protein corona enrichments. This underscores the significant potential of DISPA when paired with NP sample preparation techniques for plasma proteome studies.

## INTRODUCTION

Proteins are the molecular machines that drive cellular functions, executing most of the biological activities essential for life. The disfunction of proteins often contributes to various diseases. Alterations in the concentration or structural integrity of specific proteins can act as indicators, or biomarkers, which signal the presence of a particular disease or the physiological condition of an organism.^1, 2^ For example, cardiac troponin is a specific marker for cardiac injury ^3^ and alpha fetoprotein (AFP) levels are associated with certain types of cancer.^4^ Blood circulates throughout the body, interacting with nearly every type of cell and tissue. Consequently, various protein alterations occurring within the body can often be identified by analyzing the plasma.^5^ This has led to a rising demand for plasma proteomic analysis in clinical diagnostics. However, current plasma proteomic analysis is still hindered by the following factors: (1) there is a very high dynamic range with protein concentrations spanning more than 10 orders of magnitude for the plasma proteome.^6^ This means that highly abundant proteins can easily mask the detection of low-abundance proteins, and the latter often include important disease biomarkers.^7^ (2) Current analytical methods centered on liquid chromatography-mass spectrometry (LC-MS) produce deep coverage but are constrained by drawbacks of low throughput, high cost and inconsistent results, thereby hindering their widespread application in large cohort analysis and clinical diagnosis. ^5, 8-12^

Recently, a plethora of innovative strategies has been developed to meet these challenges. For example, techniques to remove highly abundant proteins include immunodepletion, ultrafiltration, and chromatographic fractionation to enhance the depth of plasma proteome coverage.^13^ There has also been a shift towards automation, with sample processing platforms and robotics being harnessed to standardize sample preparation.^14-16^ Furthermore, the adoption of short gradients and multiple trap columns in liquid chromatography systems is being used to maximize throughput.^17-19^ In addition to this, our previous work demonstrated a radical protocol that totally eliminates liquid chromatography,^20^ which realized a total of 2,000 protein identifications and more than 1000 protein quantifications in less than 4 minutes.^21^ Yet, a comprehensive plasma proteome analysis pipeline, which harmoniously balances all requirements including high throughput, cost effectiveness, and robustness, remains elusive.

Here, to address this bottleneck, we introduce a hybrid protocol that combines DISPA with the nanoparticle corona^22-24^ enrichment for high throughput plasma proteome analysis. We demonstrated that, in a non-targeted scenario, our method can identify up to 393 proteins in one MS injection. Our method confirms the differential protein corona profiles among various nanoparticles based on their distinct surface modifications. All nanoparticles outperformed neat plasma workflow, as evidenced by deeper coverage, more enriched biomarkers, and better quantification accuracy. Additionally, to maximize the efficiency and throughput of this protocol, we optimized it further into a targeted DISPA method, which can identify over 280 proteins, including 44 biomarkers, in just 1.4 minutes of acquisition from one particle enrichment. We present the stability, repeatability, robustness, and quantitative accuracy of this method across multiple nanoparticles and concentrations. We also designed a scoring system to evaluate the performance of individual nanoparticles or their combinations, and the carboxylic acid-modified nanoparticle ranked highest among the five tested. This study offers a novel solution for high-throughput proteomic analysis of clinical plasma samples. We are excited to further explore the potential of DISPA when integrated with specialized sample preparation techniques in clinical proteomics.

## EXPERIMENTAL SECTION

### Chemicals and standards

Angiotensin I (Sigma, A9650-1MG), QCAL Peptide Mix (Sigma, MSQC2), and HeLa digest standard (Thermo Fisher Scientific, Catalog number: 88328) were dissolved into different concentrations with 60% acetonitrile (ACN) in 0.1% formic acid (FA). Dimethyl sulfoxide (DMSO) was purchased from Sigma.

### Nanoparticles and plasma sample preparation

Proteograph™ XT Assay Kit, which contains five different nanoparticles, was used for the enrichment of plasma protein. To create the protein corona, we used the Proteograph Assay to incubate five NPs with 100 μL of plasma samples in 96-well plates using the SP100 Automation Instrument (Seer Inc.). These plates were then sealed and warmed to 37 °C for an hour while being shaken at 300 rpm. Post-incubation, a magnetic device was used to pull down the NPs from the plate for 5 minutes. The remaining liquid, which held proteins not attached to the corona, was removed. The bound protein corona was rinsed thrice using a wash buffer. It’s important to note that the samples already had EDTA as they were collected in K2 EDTA tubes. For the digestion of NP protein corona, trypsin digestion was done following the manufacturer’s instructions. Finally, peptide concentration was determined utilizing a colorimetric peptide assay kit from Thermo Fisher Scientific.

### Mass spectrometry

DISPA of the plasma proteome peptides was performed on an Orbitrap Fusion Lumos™ mass spectrometer (Thermo Fisher Scientific) coupled with the FAIMS Pro Interface. A nano-ESI source (“Nanospray Flex”) and LOTUS nESI emitters from Fossiliontech were used for ionization. The Ultimate™ 3000 HPLC system (Thermo Fisher Scientific, USA) was used to control the automated sample loading, flow rate, and mobile phase composition. A flow rate of 1.4 μL/min was maintained for the beginning 0.5 min to quickly deliver the sample to the nanoESI emitter, and then, the flow rate was reduced to 0.25 μL/min for the data collection period. An isocratic flow consisting of 60% ACN in 0.1% FA was maintained during the whole acquisition time. The performance and effectiveness of the whole system was tested with 1 fmol/μL angiotensin I prior to start any experiments and the normalized TIC signal of angiotensin I should be at least 1E5 as a baseline performance requirement. Non-targeted scouting experiments were performed using DIA across a precursor range of *m/z* 400 to 1000 with a 2 *m/z* window with stepping compensation voltages from -30V to -80V in 10V increments. Targeted DISPA analysis was conducted by only targeting specific *m/z* windows that showed peptide identifications in non-targeted DISPA.

### LC-MS analysis

Peptides were trapped on EXP®2 Stem Trap column and separated on a 200 cm µPAC™ column from PharmaFluidics. The entire assembly was connected by 20μm internal diameter Viper™ capillaries (Thermo Fisher Scientific) and kept at 55 °C in the column oven. The reversed-phase analytical gradient was delivered as follows: (mobile phase A: 0.1% formic acid in water, mobile phase B: 0.1% formic acid in acetonitrile): start at 8% B, linear ramp to 25% B at 70 min, linear ramp to 37% B at 95 min; jump to 98% B at 96 min and hold for 9 min, drop back to 8% B at 105 min and hold for 5 min (110 min total). The loading pump was connected to three solvents: 0.1% formic acid in water (loading buffer A), 0.2% formic acid in 70% acetonitrile and 30% water with 5 mM ammonium formate (loading buffer B), and 0.2% formic acid in 50% isopropanol, 30% acetonitrile, 20% water and 5 mM ammonium formate (loading buffer C). At the start of the method, the loading pump delivered 50% B and 50% C at 50 μL/min to 5 min. The solvent was switched to 100% A at 6 min and ended at 62 min. The loading pump flowrate was reduced to 10 μL/min at 70 min and held to the end of the gradient. Data dependent acquisition was applied, and tune method details set as: positive ionization (2500V), ion injection time (100ms), AGC (100%), HCD energy (30%).

### Library generation

The building of proteome library used for CsoDIAq generally included three steps: 1) performing LC-MS/MS analysis of plasma samples with data dependent acquisition (DDA) for eleven compensation voltages from -30V to -80V in a step of 5V. 2) Using Fragpipe to produce pepxml files of peptides and proteins with human fasta database (2022-08-22) and adding decoys (50%). 3) Building a library for CsoDIAq with SpectraST; for more details see supporting information.

### Identification and quantification of peptides and proteins

Peptides and proteins were identified with CsoDIAq^25^, for more detailed information about CsoDIAq see https://github.com/xomicsdatascience/zoDIAq.

### Data analysis and availability

Python version 3.9.7 and R version 4.3.0 were used for data analysis and data visualization and all code are provided open source via jymbcrc/Plasma_NP_DISPA github. All raw mass spectrometry data are deposited and available at ftp://MSV000094026@massive.ucsd.edu with password “plasmaDI”.

## RESULTS AND DISCUSSION

### Multi Nanoparticle-based sample preparation and non-targeted DISPA

**Figure 1A** depicts the workflow with the Proteograph multi-NPs enrichment assay, physicochemical attributes of NPs and direct infusion shotgun proteomic analysis (DISPA). Briefly, superparamagnetic iron oxide NPs were incubated with plasma samples to form a protein corona in an automatable Proteograph assay on the SP100 Automation Instrument. This was followed by a rapid magnetic separation (<30 seconds) from plasma, facilitated by the superparamagnetic core of the nanoparticles. The nanoparticles can accommodate diverse surface chemistries, thereby facilitating the generation of unique corona compositions that can be employed for broader proteome interrogation.^6, 23, 24^ Following the protein corona digestion, the tryptic peptides were subjected to direct infusion using a nano emitter for ionization, without the separation process of column liquid chromatography. Multiple gas-phase separation processes, including High Field Asymmetric Waveform Ion Mobility Spectrometry (FAIMS) and quadrupole selection, were employed prior to peptide entry into the Orbitrap™ MS analyzer. These separations reduce the complexity of ions and maximize peptide identifications.

**Figure 1.**
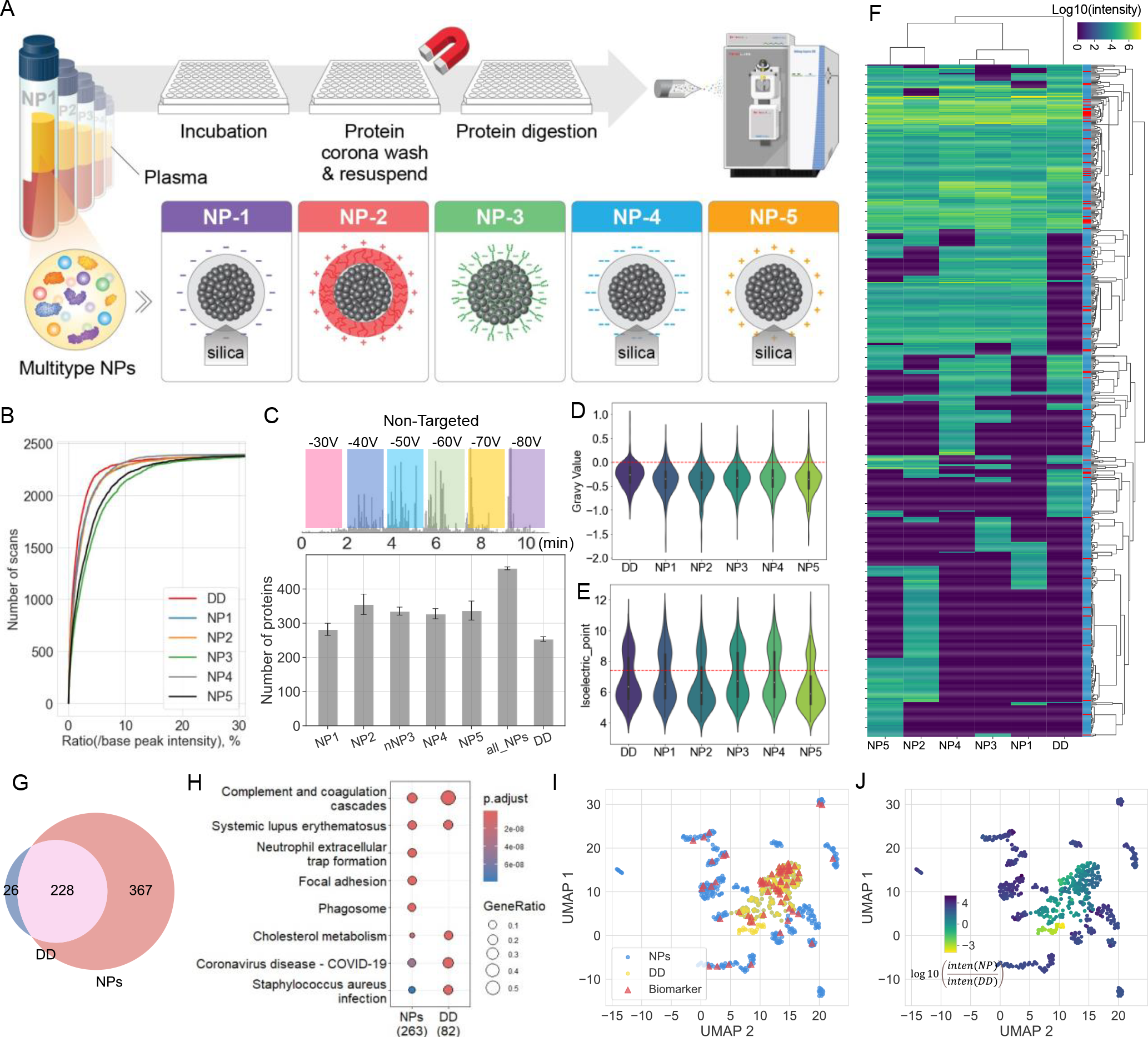
Overview of DISPA with nanoparticle enrichment. (A) workflow of multi-NP enrichment assay and physicochemical attributes of the NPs. (B) Number of scans for given intensity ratios (divided by base peak intensity) in different NPs and neat plasma. (C) Typical TIC of non-targeted DISPA analysis and bar plot showing protein identifications of different NPs (NP1-5) and direct digested neat plasma (DD) by non-targeted DISPA method (n=3). (D) GRAVY Values of all identified proteins for NPs and neat plasma. (E) Isoelectric points of all identified proteins from NPs and neat plasma. (F) Heatmap showing the protein intensities identified by DISPA across 5 NPs and neat plasma with biomarkers in red along the right. (G) Venn diagram showing the protein groups overlap between neat plasma and NPs using non-targeted DISPA. (H) KEGG pathway enrichment analysis of NPs and neat plasma using Fisher’s exact test and p-value <0.05 corrected by Benjamini–Hochberg. GeneRatio indicates genes in each group to the total significant Genes. (I, J) Dimension reduction of the protein profiles into 2D using UMAP comparing direct digested neat plasma (DD) and NPs. Each dot represents the intensities of a given protein group across all NPs and neat plasma. The color of the dot denotes the class of corresponding protein profile (I), or the log10-fold change of a given protein intensity between Direct Digested neat plasma and NPs (J). Plasma biomarkers are highlighted by red stars. Undetected proteins in any NPs or neat plasma are assigned an intensity value of 10.

The five distinct types of nanoparticles (NPs) were tested for their efficacy, initially through non-targeted DISPA scouting experiments. These experiments involved scanning from *m/z* 400 to 1000 with a DIA window of 2 Da for each compensation voltage from -30V to -80V in a step of 10V (**Figure S1**). The Total Ion Chromatogram (TIC) indicated that typical NP-enriched samples manifested less low intensity scans (below 10% in comparison to base peak intensity) than neat plasma samples that had undergone direct digestion (**Figure 1B, Figure S2**), which suggests a diminished ion competition effect typically associated with high abundance proteins, enabling successful identification of low abundance proteins along with high abundance proteins without a need for depletion. In agreement with this, we further observed a higher number of peptide and protein identifications in NP-processed samples compared to the neat plasma digestion workflow using non-targeted DISPA (**Figure 1C, Figure S3**). Notably, an average of over 300 protein identifications in replicate MS runs was achieved by 4 out of the 5 NPs, with NP2 displaying the highest, averaging over 350 protein identifications and a maximum of 393 protein groups.

Assessment of the GRAVY (Grand Average of Hydropathy) values corresponding to the identified proteins by different NPs indicated that the majority exhibit hydrophilic characteristics (**Figure 1D**), which is consistent with the characteristics of plasma proteomics. Additionally, a comparative evaluation of the isoelectric point distributions revealed slight differences among the distinct NPs and neat plasma (**Figure 1E**). These variations can be attributed to differential surface modifications and charge states of the respective NPs. For example, polymer modified NP2 and silica modified NP4, which exhibited a positive surface zeta potential, enriched more proteins with isoelectric points less than 7.4, as those proteins are negatively charged at this pH value. Further, we analyzed the unique preference between NPs and neat plasma samples.

**Figure 1F** summarizes the 621 identified proteins and their corresponding intensity across all NP coronas and neat plasma (**Supplementary Table 1**). The results indicate that while there are proteins shared across all treatments, each NP displays a distinct profile. Intriguingly, the neat plasma appears to be nearly a subset of the NPs group except for 26 unique proteins (**Figure 1G, Figure S4**). Importantly, upon comparing the identified protein groups with the 109 FDA-approved biomarkers^26^, our findings reveal that a total of 52 biomarkers were identified by NPs, with NP2 alone accounting for a maximum of 46 biomarkers (**Figure S5**). KEGG and GO pathway enrichment analyses further substantiated the broader coverage for NP coronas with more proteins exhibited enhanced enrichment in pathways, including phagosome, focal adhesion, and neutrophil extracellular trap formation etc. (**Figure 1H, Figure S6, S7**). UMAP was used to describe the quantitative profile of each protein across clusters, and whether each protein was found in NPs or neat plasma. When overlaid with biomarker annotations (**Figure 1I**), it revealed that many clusters of proteins found only with NPs contribute additional biomarkers, which is also seen by their increased abundance (**Figure 1J**).

### Targeted DISPA of plasma proteome derived from multi-NPs

In non-targeted DISPA analysis, each DIA window within a specified *m/z* range is systematically scanned across all compensation voltages. This methodology frequently results in an abundance of scans with no peptide identifications, consequently prolonging the collection time with no benefit. For high throughput DISPA, it’s customary to ascertain DIA windows enriched with peptides after the initial non-targeted approach. We used the identifications from the DISPA scouting experiments to design short, targeted injection sequences. **Figure 2A** demonstrates a perfectly overlapping total ion chromatograms (TIC) from three replicates of a typical targeted DISPA conducted on the plasma proteome originating from NP_2_ corona. In this example, the acquisition time is shortened to 1.4 min, which represents a possible throughput of over 1,000 samples per day. Importantly, while the identified protein groups diminished in comparison to non-targeted methodologies, over 280 proteins were nonetheless identified by targeted DISPA for NP_2_ and NP_5_, which equates to a rate of more than 3 proteins per second (**Figure 2B**). A total of 405 unique protein groups were identified using separate data from all NPs. A positive correlation between sample concentration and the number of identified peptides and proteins was observed in all NPs and neat plasma (**Figure 2B, Figure S8**). Specifically, with a reduction in sample concentration (after enrichment), transitioning from 0.4 µg/µl to 0.05 µg/µl, there was an approximate decrement of 10-20% in peptide and protein identifications. The reproducibility, repeatability, and robustness of targeted DISPA is demonstrated by a Venn diagram of protein and peptide identifications (**Figure 2C, Figure S9, S10**) and correlation analysis across different concentrations (**Figure 2D-E, Figure S11**) among three replicates.

**Figure 2.**
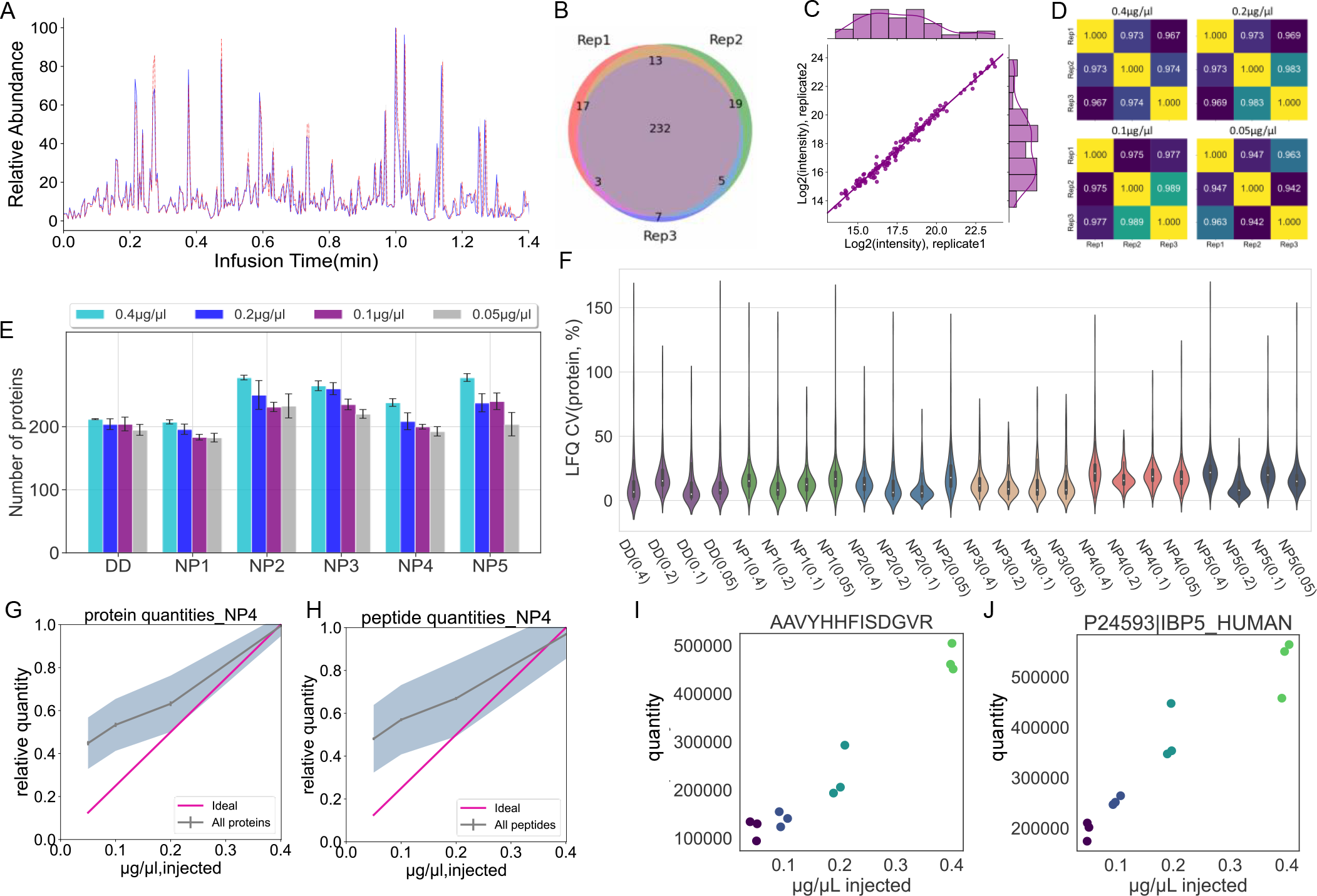
Reproducibility and quantification performance using DISPA and NP enrichment. **(A)** TIC overlay of three injections of targeted DISPA analysis. **(B)** Venn diagram of protein group overlap among three injections. Concentration:0.2 µg/µl. **(C)** Scatterplot of quantified proteins in two injections of targeted DISPA (R square = 0.988). **(D)** correlation between replicates across four different concentrations of a typical NP (NP_3_). **(E)** Bar plot showing identified protein groups of different NPs and direct digested neat plasma (DD) by targeted DISPA across four different concentrations (n=3). **(F)** Coefficient of variation for quantified proteins in four concentrations across each NP and neat plasma. The average CV among all concentrations and NPs are 17.5%. **(G)** Protein and **(H)** peptide quantities determined by targeted DISPA-LFQ from a dilution series of plasma proteome extracted with NP4. The shaded area represents one standard deviation from the mean in the middle. Protein groups were filtered for complete identifications across three replicates (n=3). **(I-J)** Examples of one quantified peptide **(I)** and protein **(J)**.

Beyond the quantity of identified peptides and proteins, the precise quantification is critical. We applied our previously reported label-free quantification (LFQ) strategy^21^ based on fragment ion intensities for the accurate quantification of plasma proteome. Firstly, the coefficient of variation (CV) of peptides and proteins (17.5% and 18.1%, respectively) derived from all specimens, including various NPs and neat plasma over four concentration levels, underscores excellent precision and robustness of the quantification methodology for plasma proteome analysis (**Figure 2F, Figure S12**). Moreover, using NP_4_ as a representative, 127 out of 149 shared proteins (identified in all concentrations and replicates) demonstrate positive slope (linear regression) and p-values (non-correlation test) less than 0.05 from Pearson correlation. This represents over 85% of the repeatably identified proteins (**Figure 2G**). These shared proteins correspond to 612 peptides, of which 438 exhibited good linearity with positive slopes and p-values less than 0.05 from Pearson correlations (**Figure 2H**). Examples of a typical quantified peptide and protein from NP_4_ are shown in **Figures 2I** and **2J**, respectively. The LFQ DISPA for plasma proteome derived from other NP coronas demonstrated same performance, with most peptides and proteins show positive slope and p-values less than 0.05 from Pearson correlation (**Figure S13**).

Next, we further explored the performance and protein functions of targeted DIPSA together with each NP and neat plasma. **Figure 3A** shows targeted DISPA intensity profiles for 443 individual protein groups that were filtered for complete identifications across three replicates in each condition (**Supplementary Table 2, Figure S14**). Interestingly, NP coronas each identified more proteins than neat plasma and each NP reveals unique protein groups. High intensity proteins are more overlapped among all sample preparation conditions. Further KEGG and GO pathway enrichment analysis proved that NPs tend to enrich more proteins in each pathway but demonstrated stronger effects on Focal adhesion and ECM-receptor interaction (**Figure 3B, Figure S15**). Details of preference of GO and KEGG enrichment analysis for each NP corona and neat plasma were shown in **Figure S16-17**. To better illustrate the difference among multiple NPs and neat plasma, UMAP was employed to compare the intensity profiles of each protein group across all NPs and neat plasma. We observed clusters that are specific to direct digestion, nanoparticles, or both, which reflect the significant variances for each protein group across multiple NPs and neat plasma (**Figure 3C**). Such differences can be attributed to the various surface modifications of NPs. Remarkably, all FDA-approved biomarkers are encompassed by the NPs, and a majority display greater intensity compared to neat plasma (**Figure 3C, D**). The dynamic intensity range of all proteins detected in each NP corona and neat plasma, as well as the quantity of enriched biomarkers, is presented in **Figure 3E**. Notably, NP_2_ alone enriches 39 biomarkers and performs best among all NPs. **Figure 3F** further demonstrates the associated UniProt IDs and classifications for all detected FDA-approved biomarkers.

**Figure 3.**
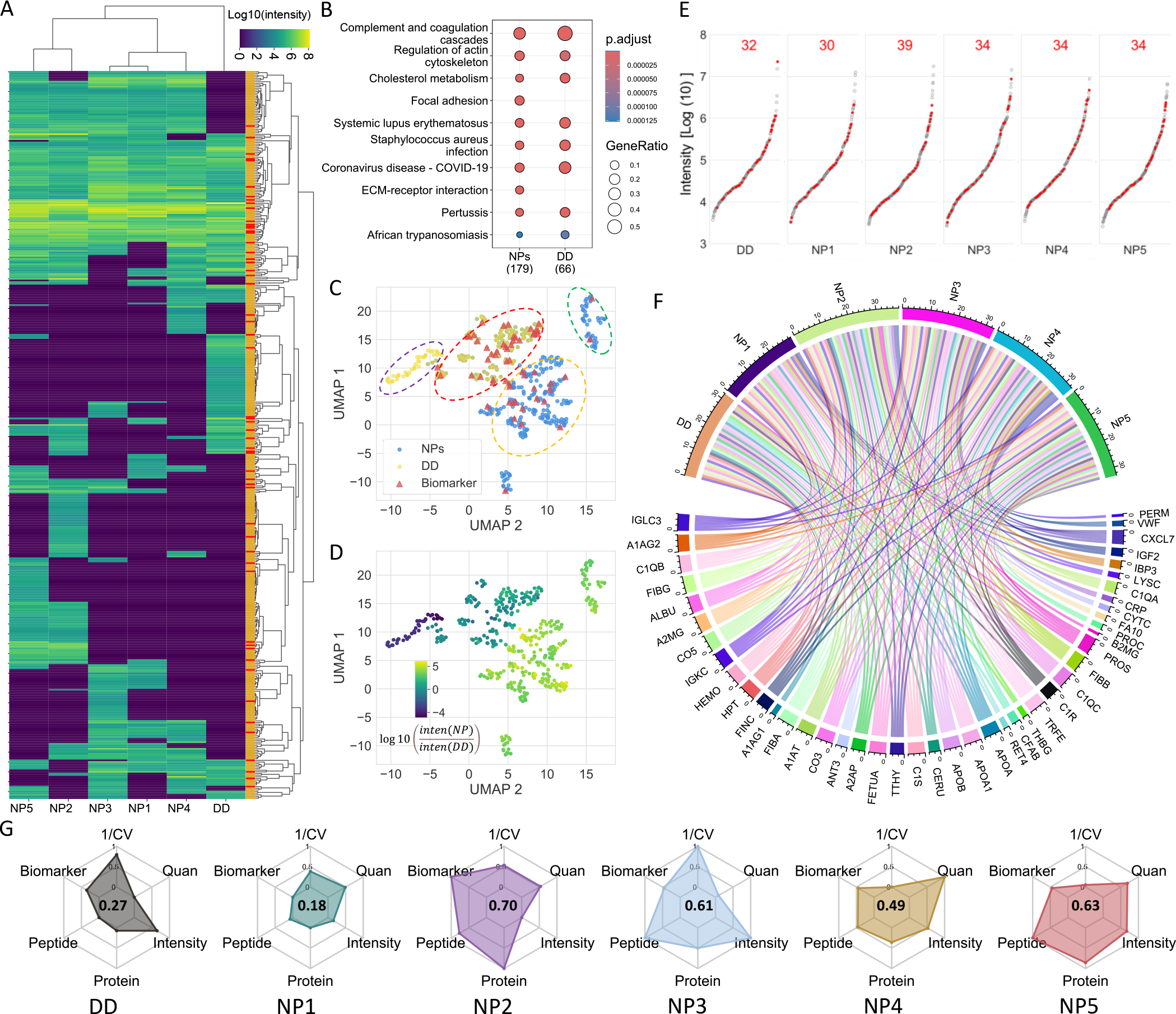
Biological Summary of Targeted DISPA results across NPs. **(A)** Heatmap showing the protein intensities identified by targeted DISPA method across 5 NPs and DD with, biomarkers noted in red along the right side. Protein groups were filtered for complete identifications across three replicates (n=3). **(B)** KEGG pathway enrichment analysis of NPs and neat plasma using Fisher’s exact test and p-value <0.05 corrected by Benjamini–Hochberg. **(C, D)** Dimension reduction of protein profiles into 2D using UMAP to compare DD and NPs. Each dot represents the intensities of a given protein group across all NPs and neat plasma. The color of the dot denotes the class of corresponding protein profile **(C)**, or the log10-fold change of a given protein intensity between Direct Digested neat plasma and NPs **(D)**. Plasma biomarkers are highlighted by red stars. Undetected proteins in any NPs or neat plasma are assigned an intensity value of 1. Protein groups were filtered for complete identifications across three replicates (n=3). **(E)** The intensity dynamic range of plasma proteome detected by targeted DISPA for all NPs and neat plasma with FDA-approved biomarkers highlighted in red and count of identified biomarkers annotated on the top. **(F)** Chord diagram showing overlap of NPs and each biomarker’s gene name. **(G)** Radar chart demonstrating the overall performance and score of each NP and neat plasma with features including peptide coverage, protein coverage, biomarkers, mean intensity, quantification performance and reciprocal of CV.

Finally, we evaluated the performance of various NPs for plasma proteome analysis when integrated with the DISPA methodology. We summarized six features for assessing the performance of these NPs, namely, peptide identification, protein identification, biomarker enrichment, quantitative precision, robustness, and intensity. Based on their importance, we weighted each feature for the aggregated score:

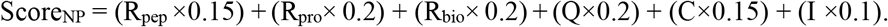

Here, “R_pep_” represents the ratio of peptides identified by NP divided by the size of peptides library (total number of peptides identified by all NPs and neat plasma together). “R_pro_” stands for the ratio of NP-identified proteins divided by proteins library size, and “R_bio_” indicates the ratio of NP-detected biomarkers divided by biomarker library size. “Q” denotes the quantification accuracy, calculated as the percentage of proteins with positive slope and p-values less than 0.05 from Pearson correlation. “C” represents robustness, which is the reciprocal of coefficient of variation, and “I” is the intensity, calculated as the mean intensity of all identified proteins.

The scores of multiple NPs and neat plasma on these six features are delineated in **Figure 3F**, with NP_2_ delivering the highest aggregated score. Meanwhile, considering the observed overlaps and disparities among the proteomes enriched by distinct nanoparticles (**Figure S18**), we conducted a comprehensive assessment of various combinations. Our findings demonstrate that, when only two NPs can be chosen, the synergistic effect of NP_2_ and NP_3_ is optimal (**Figure S19**). Among the combinations of 3 NPs, the trio of NP_2_, NP_3_, and NP_4_ yielded the highest efficacy (**Figure S20**). Notably, for configurations allowing a quartet choice, the ensemble of NP_2_, NP_3_, NP_4_, and neat plasma exhibited superior performance (**Figure S21**).

## CONCLUSIONS

In conclusion, we introduce a streamlined protocol for plasma proteome analysis that is not only high-throughput but also time-efficient, cost-effective, straightforward, and user friendly. This method integrates DISPA with NP protein corona dynamic range compression to achieve both higher speed and deeper plasma proteome coverage. The performance of this method is now optimized to the rate of over 280 protein identifications in 1.4 minutes, and 405 unique protein groups with all 5 NPs combined in approximately 7 total minutes of data collection. This approach is attractive due to the sensitivity, robustness, and diversity of corona proteomes delivered by different nanoparticles. This study presents an alternative approach to high-throughput plasma proteome analysis by replacing liquid phase separation with gas phase separation to enhance analytical throughput. We are optimistic about the expanding applications of DISPA in clinical diagnostics, especially when paired with complementary techniques like NP enrichment.

## ACKNOWLEDGMENTS

This work was supported by the United States National Institute of Health (NIH) NIGMS R35 GM142502.

