## Supplemental methods and figures for "1.4 min Plasma Proteome Profiling via Nanoparticle Protein Corona and Direct Infusion Mass Spectrometry"

### Table of contents

#### Supplemental Methods

##### Identification and quantification of peptides and proteins

CsoDIAq is a python software package designed to enhance usability and sensitivity of the projected spectrum−spectrum match scoring concept. The process was described as the following steps: Firstly, the original Thermo Fisher Scientific .RAW files were converted to mzXML files using msconvert with default settings. Then the output mzXML files were input to CsoDIAq GUI and the appropriate spectral library for that sample was selected with the default settings keeping the fragment mass tolerance at 20ppm and the "Label free quantification" should be selected. CsoDIAq produces three output files for each input mzXML file that report spectra, peptides, and proteins filtered to <1% FDR. In each case, CsoDIAq sorts peptide identifications by match count and cosine (MaCC) score, calculates the FDR for each identification using a modification of the target-decoy approach where FDR at score S = number of decoys/number of targets, and removes SSMs below a 0.01 FDR threshold. The peptide FDR calculations only use the highest-scoring instance among all SSMs for each peptide. CsoDIAq uses the IDPicker algorithm to identify protein groups from the list of discovered peptides and adds them as an additional column in the output. A detailed description about data processing, FDR calculation and protein inference was listed in our previous paper (Cranney *et al.*, Analytical Chemistry, 2021). Peptides and proteins were quantified using CsoDIAq to extract the sum of all detected fragment ion intensities of common peptides in all input files.

#### Supplemental Figures

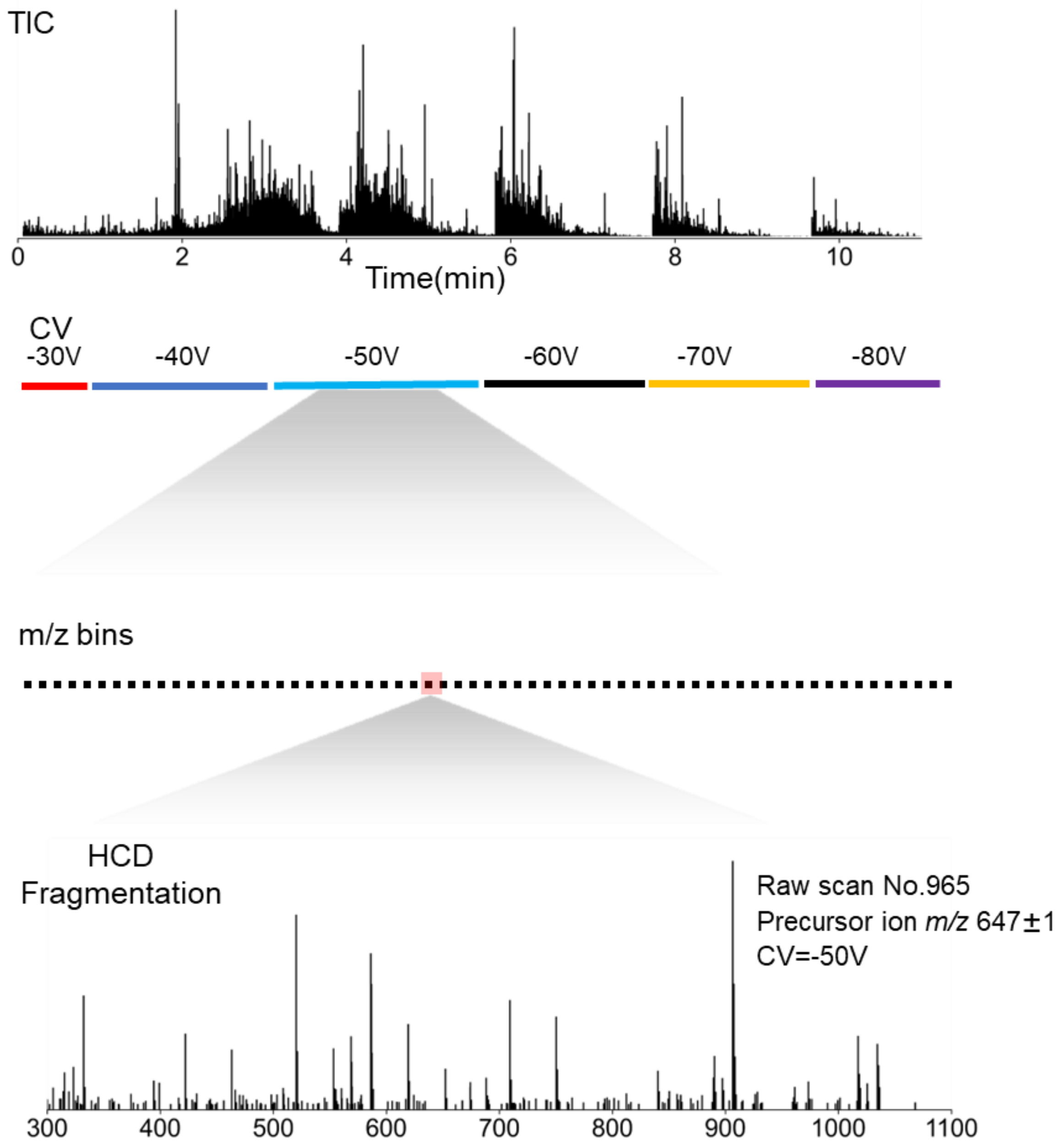


**Figure S1.** Schematic diagram of a representative targeted experiment with actual data from NP3-. Peptides were directly infused into the mass spectrometer over the duration of the experiment. Six FAIMS CVs were selected sequentially, and within each CV, specific *m/z* regions were selected with Q1 before fragmentation to produce chimeric MS/MS spectra.

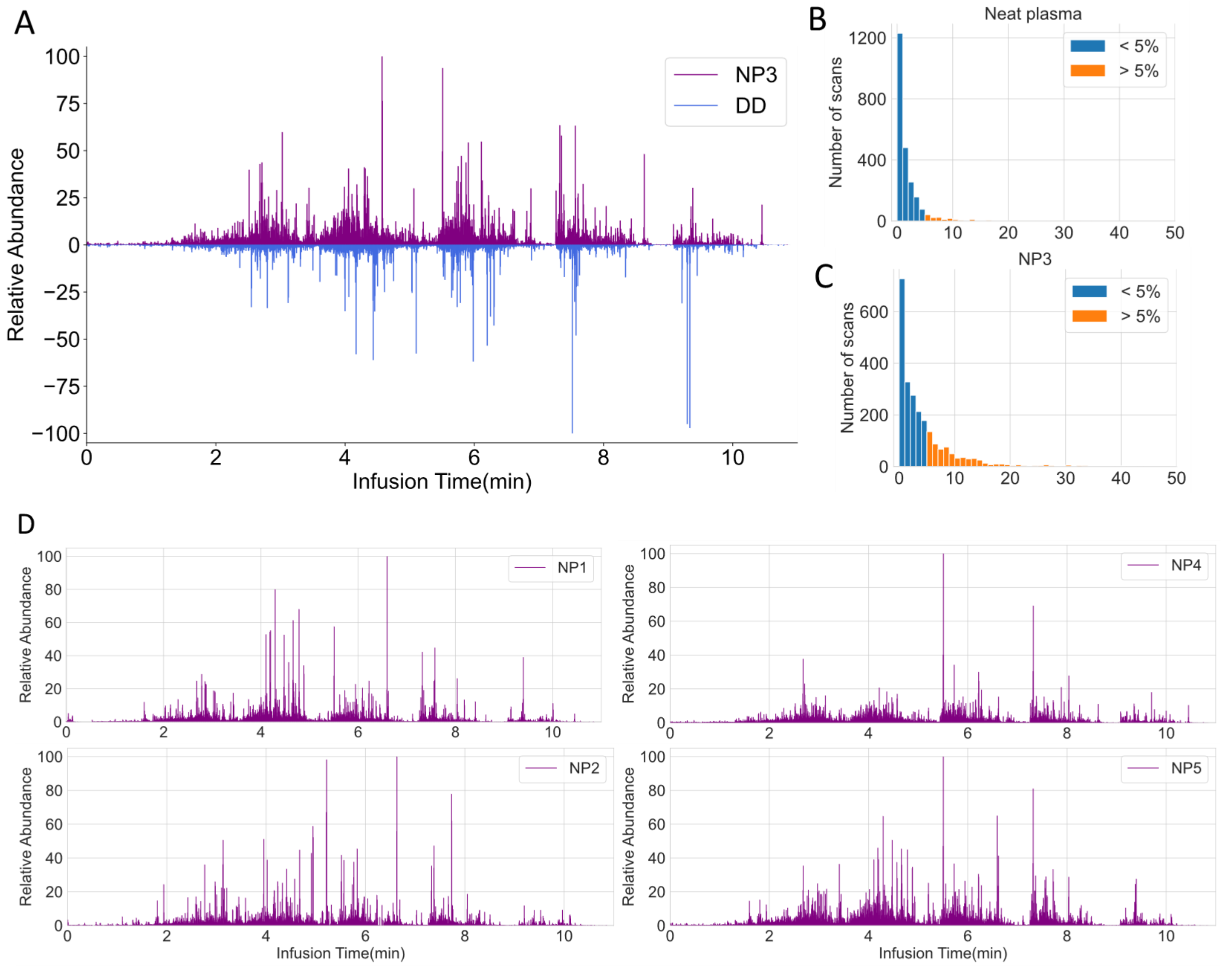


**Figure S2.** (A) Comparison of TIC derived from non-targeted DISPA between a typical nanoparticle (NP3) and neat plasma processed peptide chromatogram profile. (B-C) Histogram of intensity ratios for all scans compared to base peak in neat plasma (B) and NP3 (C), respectively. (D) Typical TIC of non-targeted DISPA for different NPs.

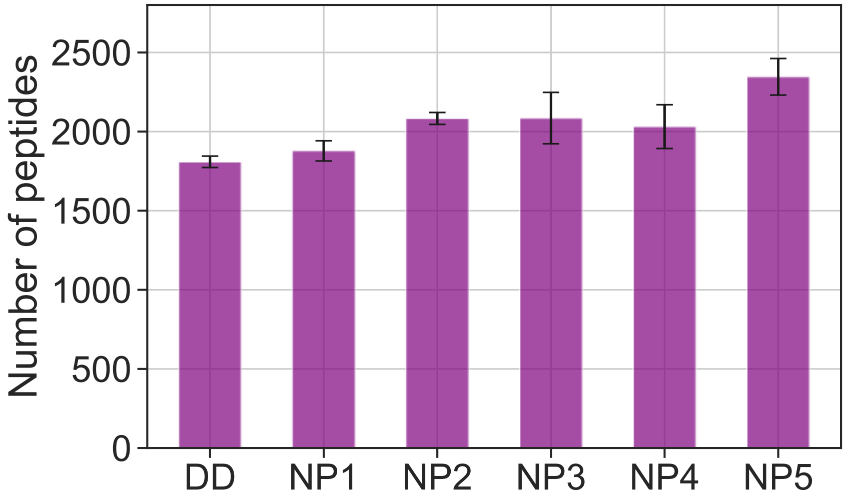


**Figure S3.** Bar plot showing peptide identifications among different NPs (NP1-5) and neat plasma (DD).

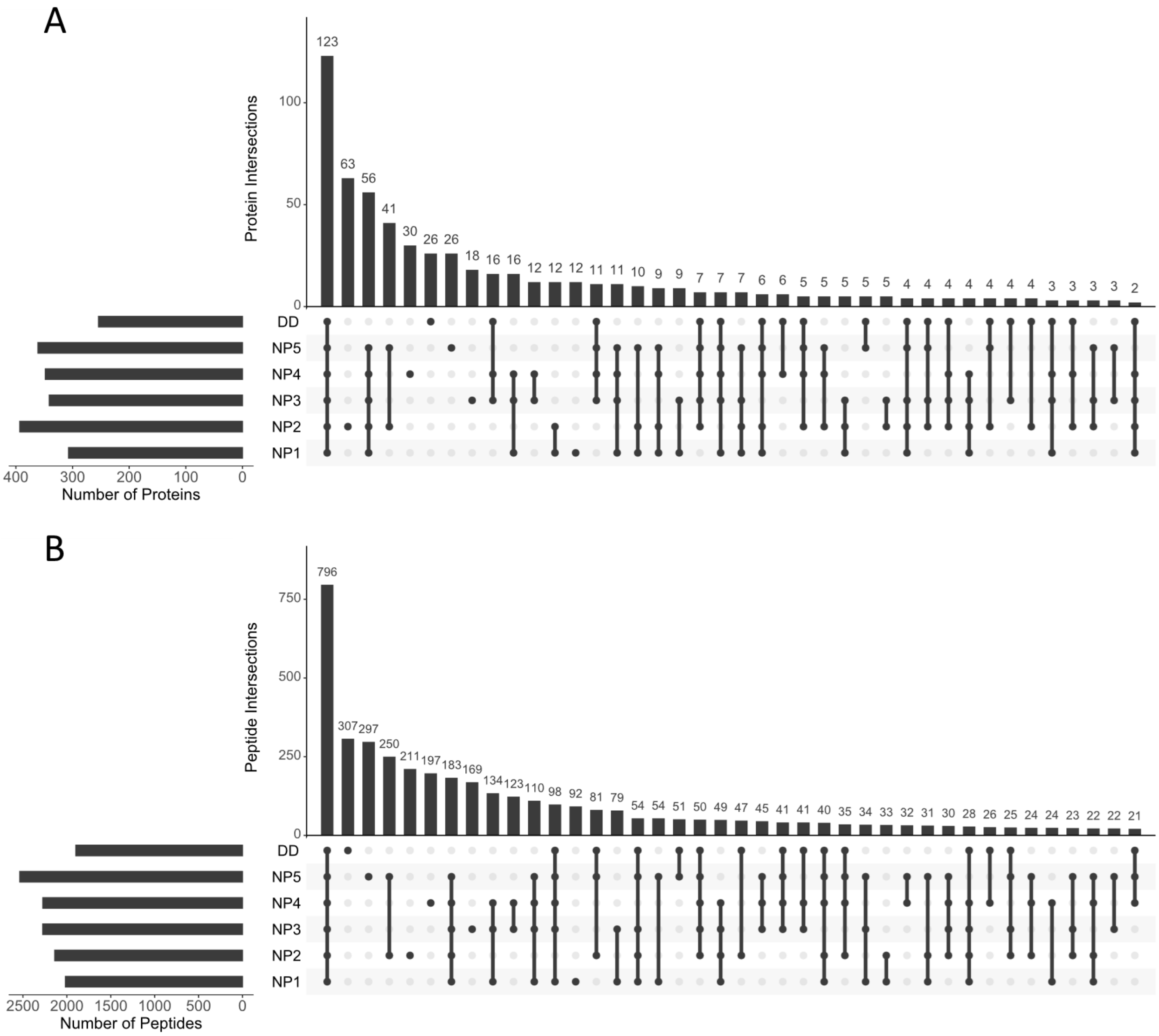


**Figure S4.** UpSet plot showing the overlap of identified proteins (A) and peptides (B) among 5 NPs and neat plasma by non-targeted DISPA.

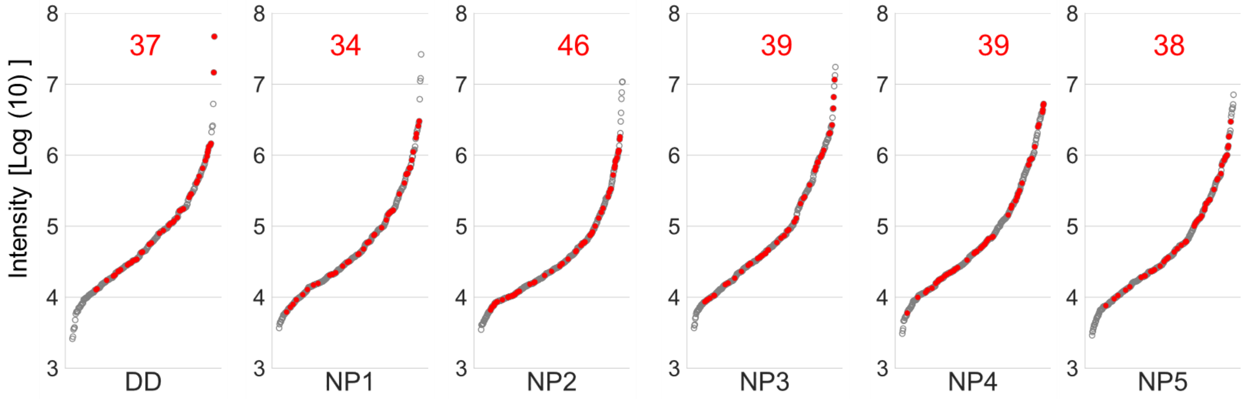


**Figure S5.** The intensity dynamic range of plasma proteome detected by non-targeted DISPA for all NPs and neat plasma, FDA-approved biomarkers were highlighted in red and quantity of identified biomarkers annotated on the top.

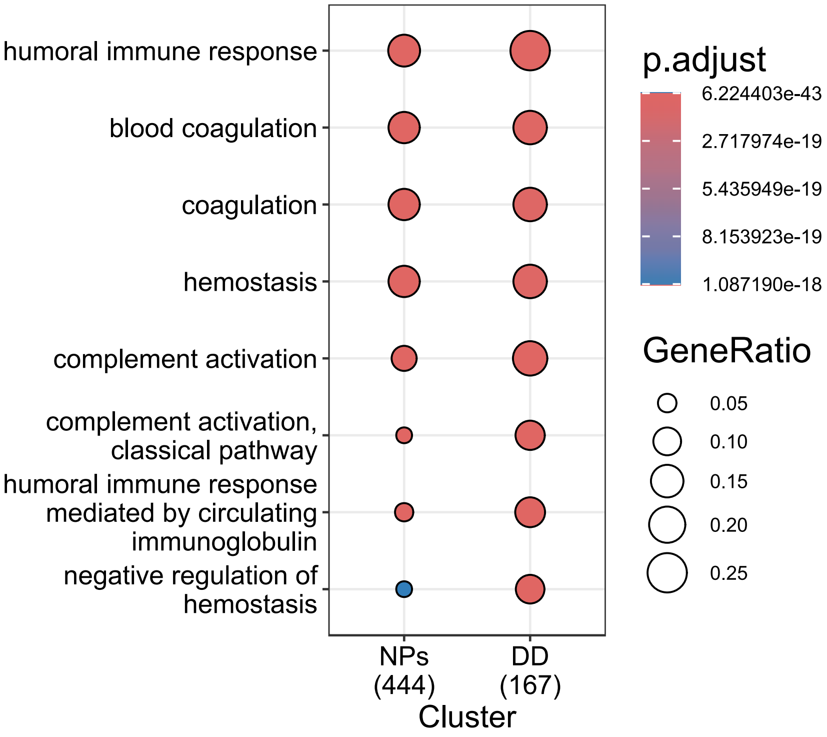


**Figure S6.** GO pathway enrichment analysis of NPs and neat plasma using non-targeted DISPA. Fisher's exact test was applied and p-value <0.05 corrected by Benjamini–Hochberg.

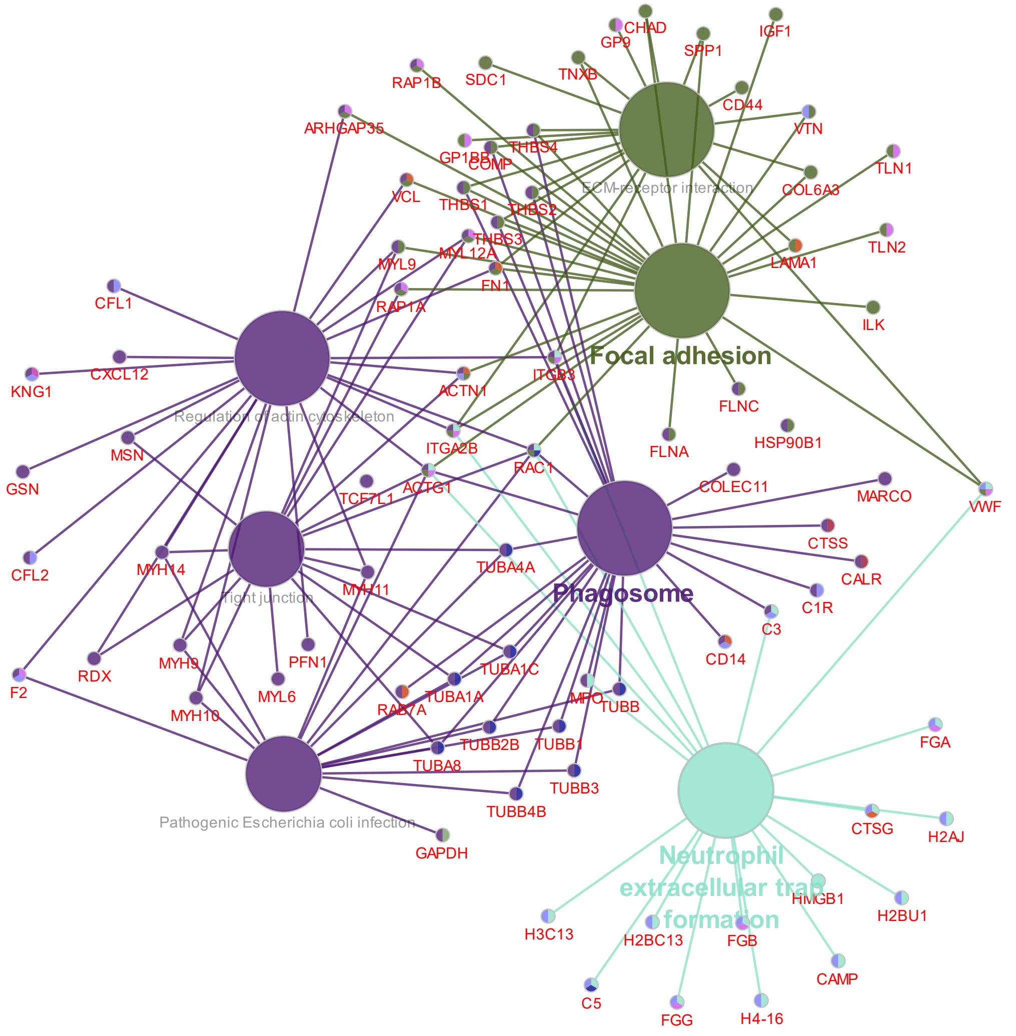


**Figure S7.** Protein members of pathways from KEGG enrichment analysis with unique proteins identified by nanoparticles. P-value cutoff set at 0.05 for term enrichment.

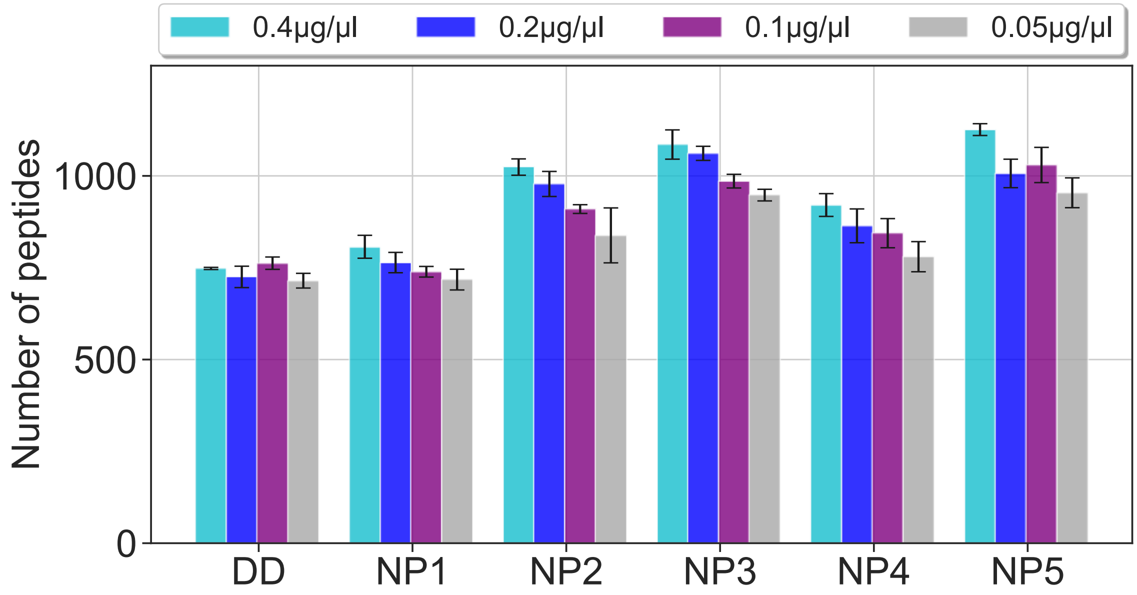


**Figure S8.** Bar plot showing identified peptides of different NPs and direct digested neat plasma (DD) by targeted DISPA across four different concentrations (n=3, concentration: 0.4 µg/µl, 0.2 µg/µl, 0.1µg/µl, 0.05 µg/µl,).

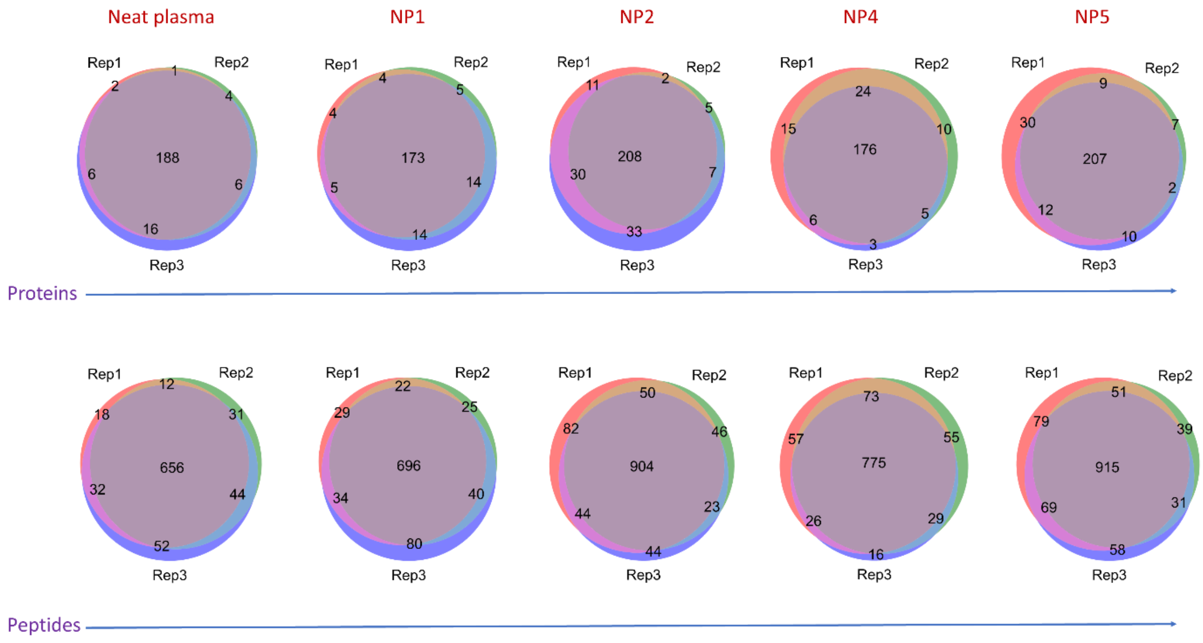


**Figure S9.** Venn diagram demonstrates the protein and peptide overlap among three injections in various NPs and neat plasma (DD) by targeted DISPA.

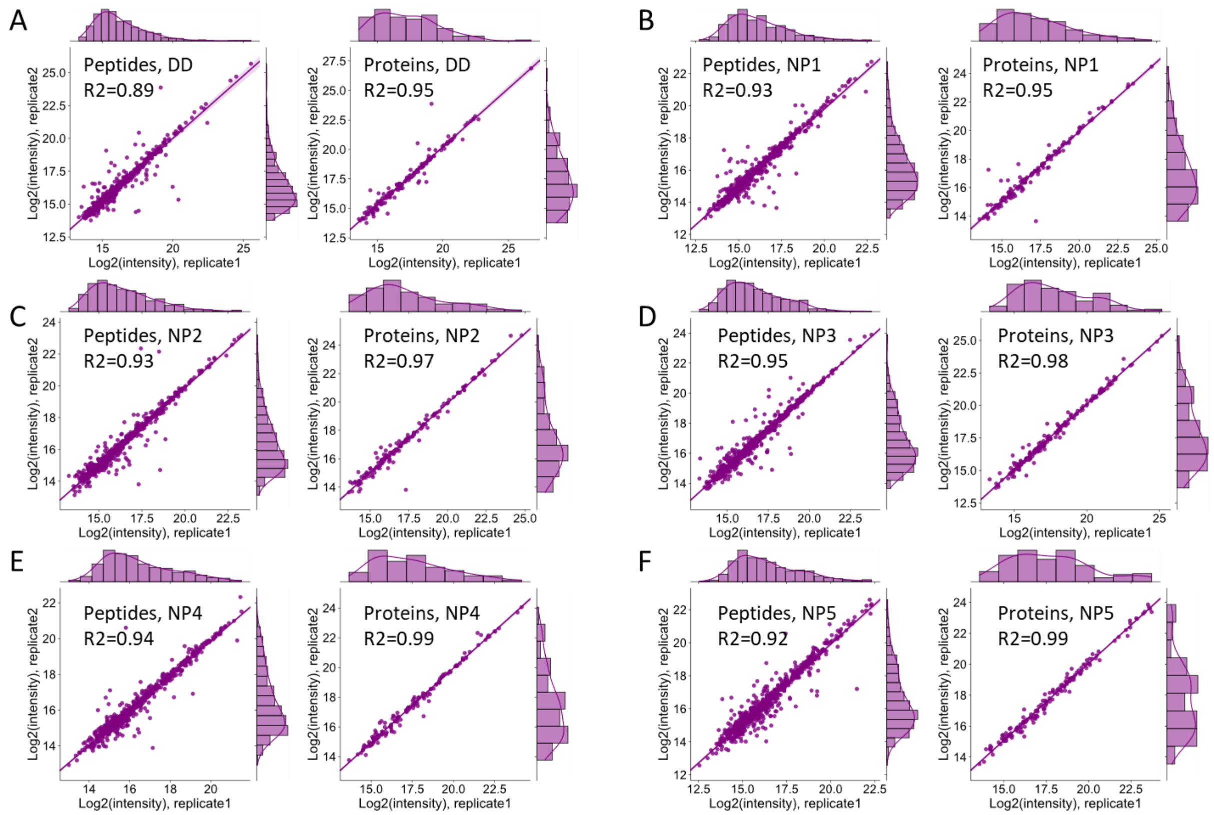


**Figure S10.** Scatterplot showing the reproducibility of all identified peptides and proteins in neat plasma (A) and different NPs (B-F).

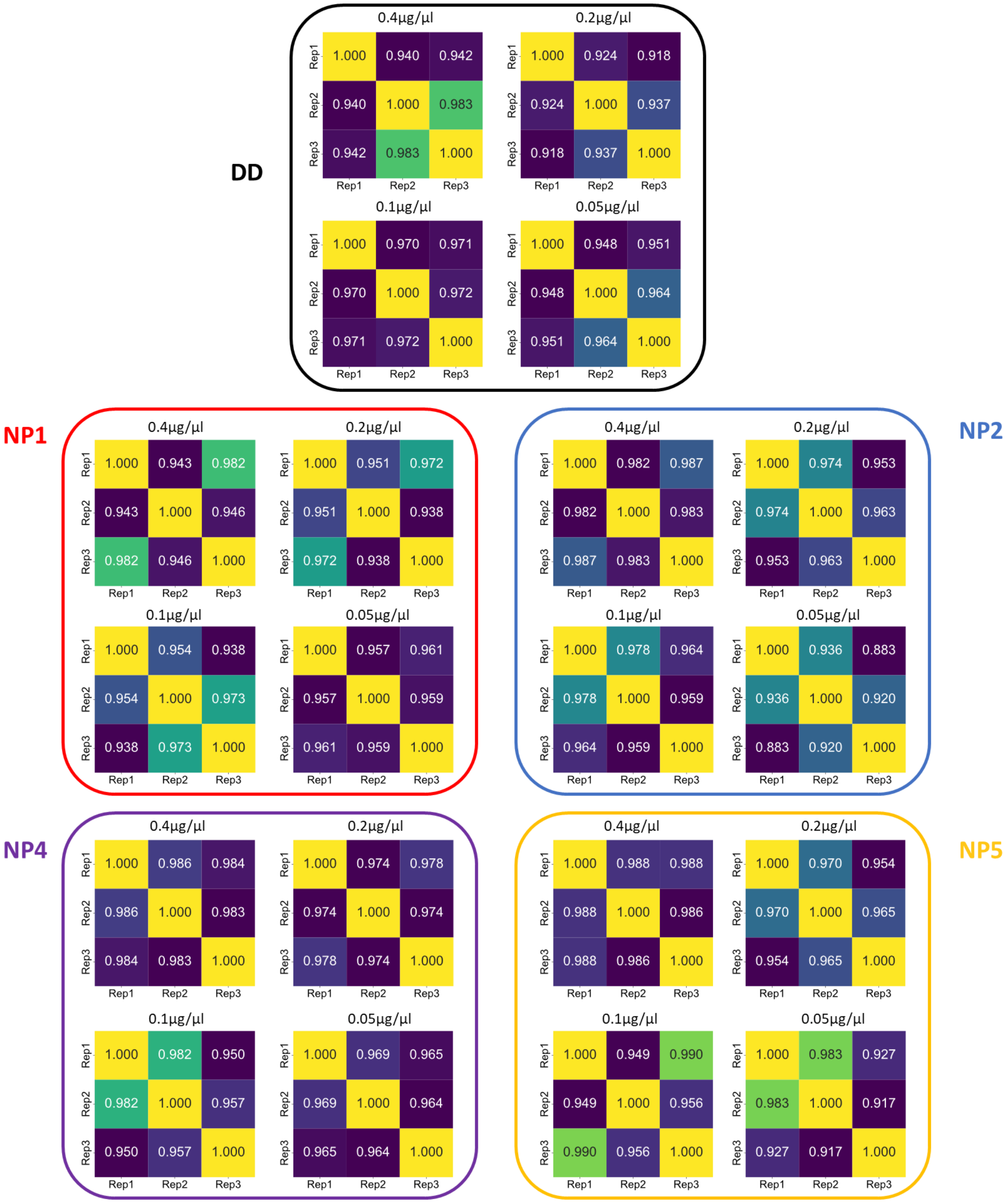


**Figure S11.** Correlation between replicates across four different concentrations of direct digested neat plasma (DD) and various NPs (NP1, 2, 4, 5).

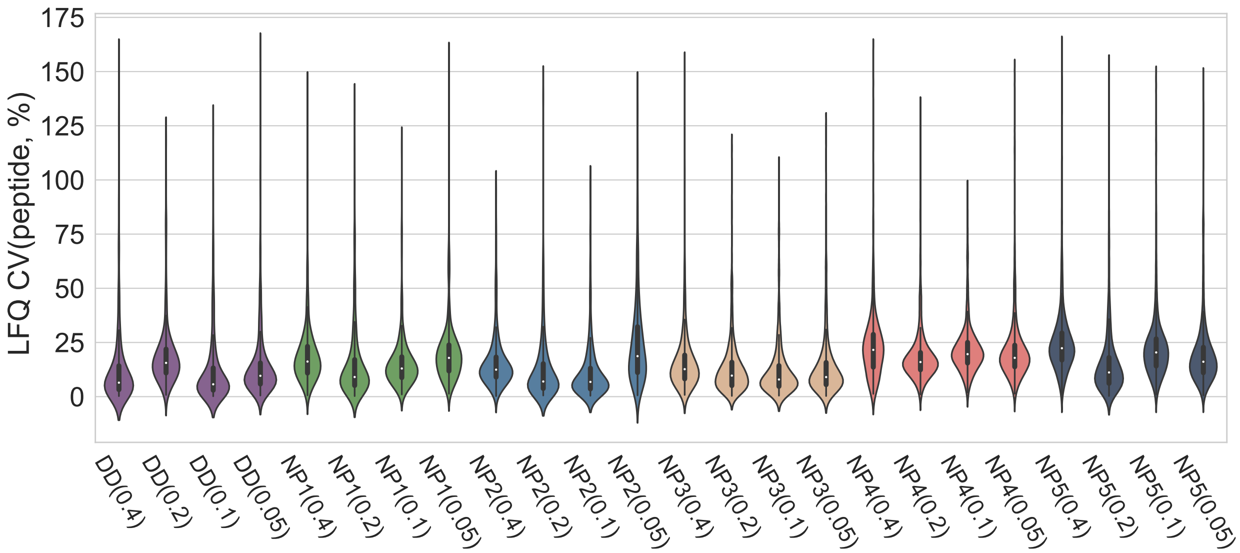


**Figure S12.** Coefficient of variations for quantified peptides within each NP and concentration. (The average CV among all concentrations and NPs are 18.1%). DD: direct digested neat plasma.

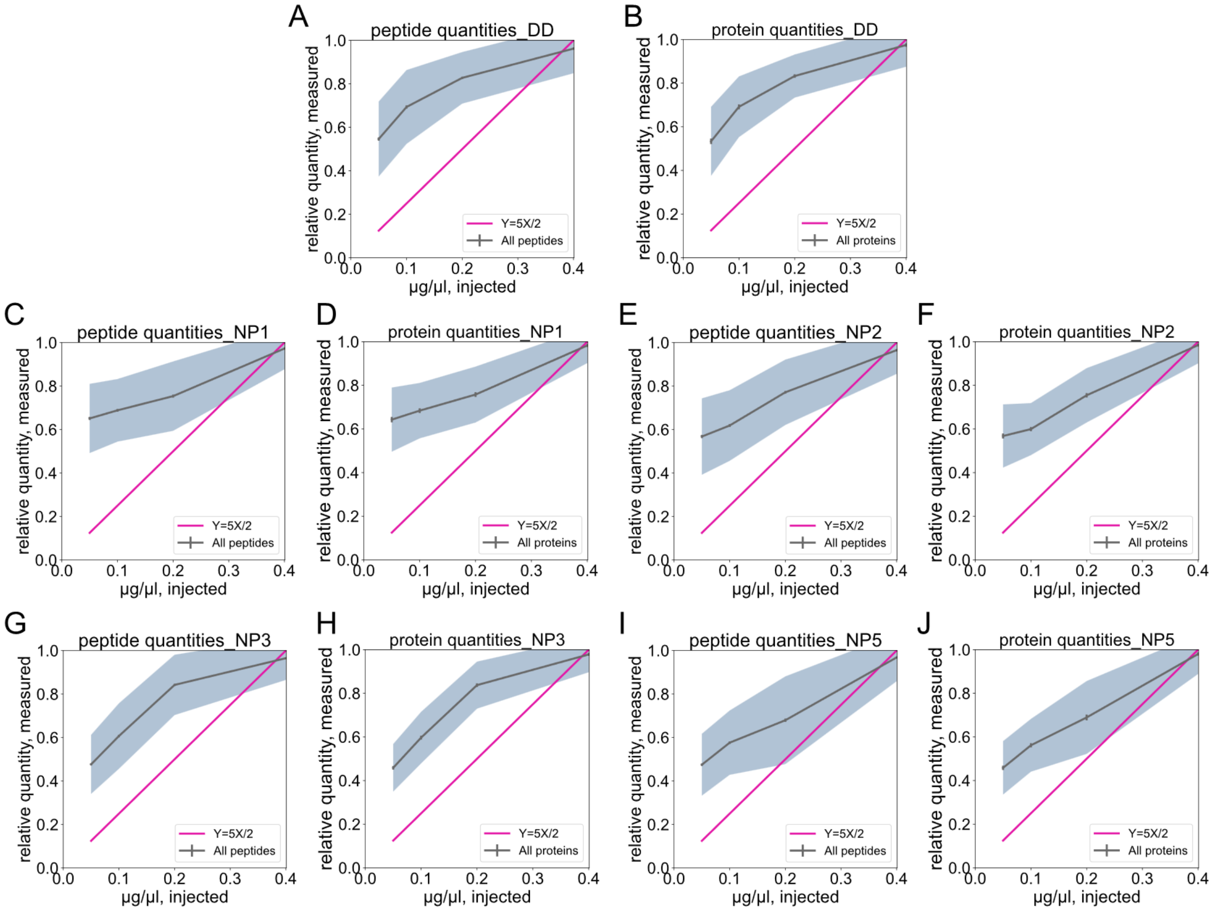


**Figure S13.** Protein and peptide quantities determined by targeted DISPA-LFQ from a dilution series of plasma proteome derived from direct digested neat plasma (A, B), NP1 (C, D), NP2 (E, F), NP3 (G, H) and NP4 (I, J). The shaded area represents one standard deviation from the mean in the middle.

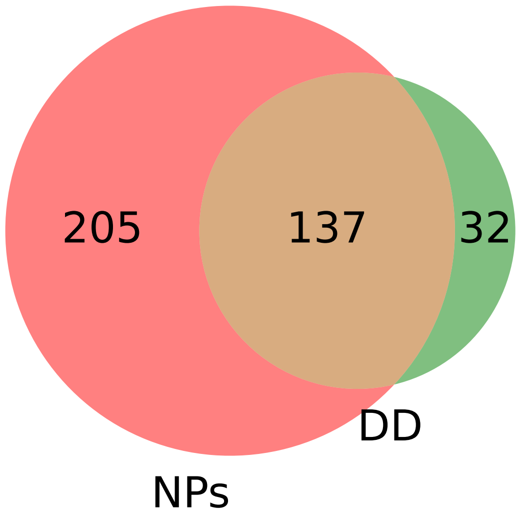


**Figure S14.** Venn diagram demonstrates overlap of protein identification between NPs and DD with targeted DISPA method.

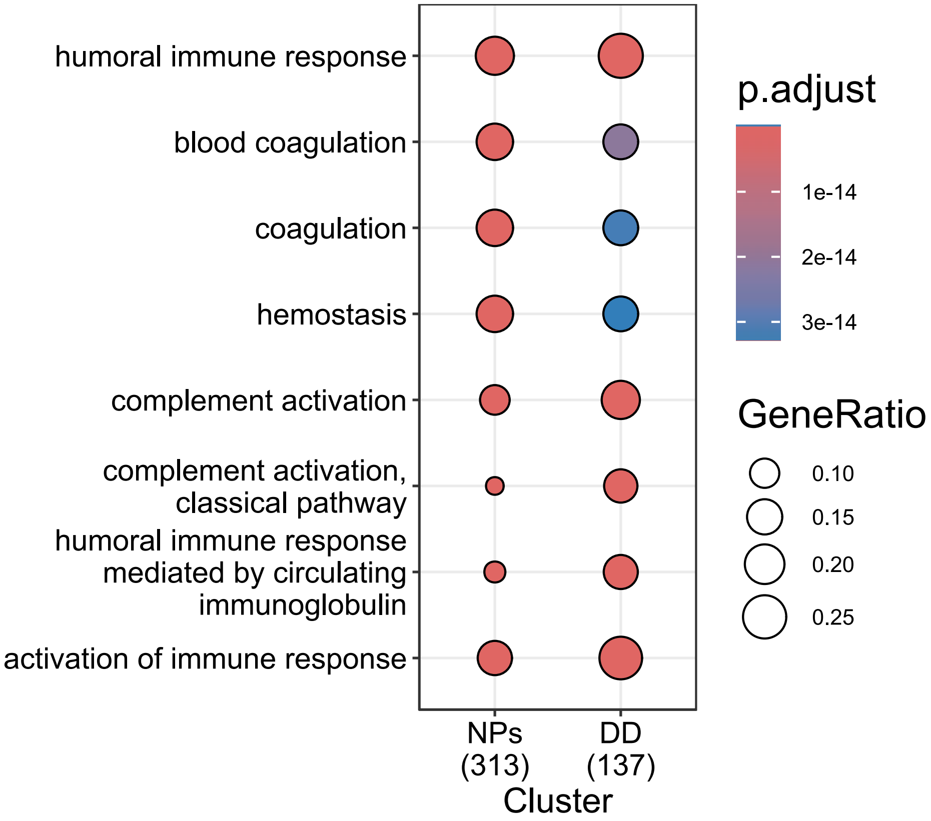


**Figure S15.** GO pathway enrichment analysis between all NPs and neat plasma (DD) by targeted DISPA. Fisher's exact test was applied and p-value <0.05 corrected by Benjamini–Hochberg.

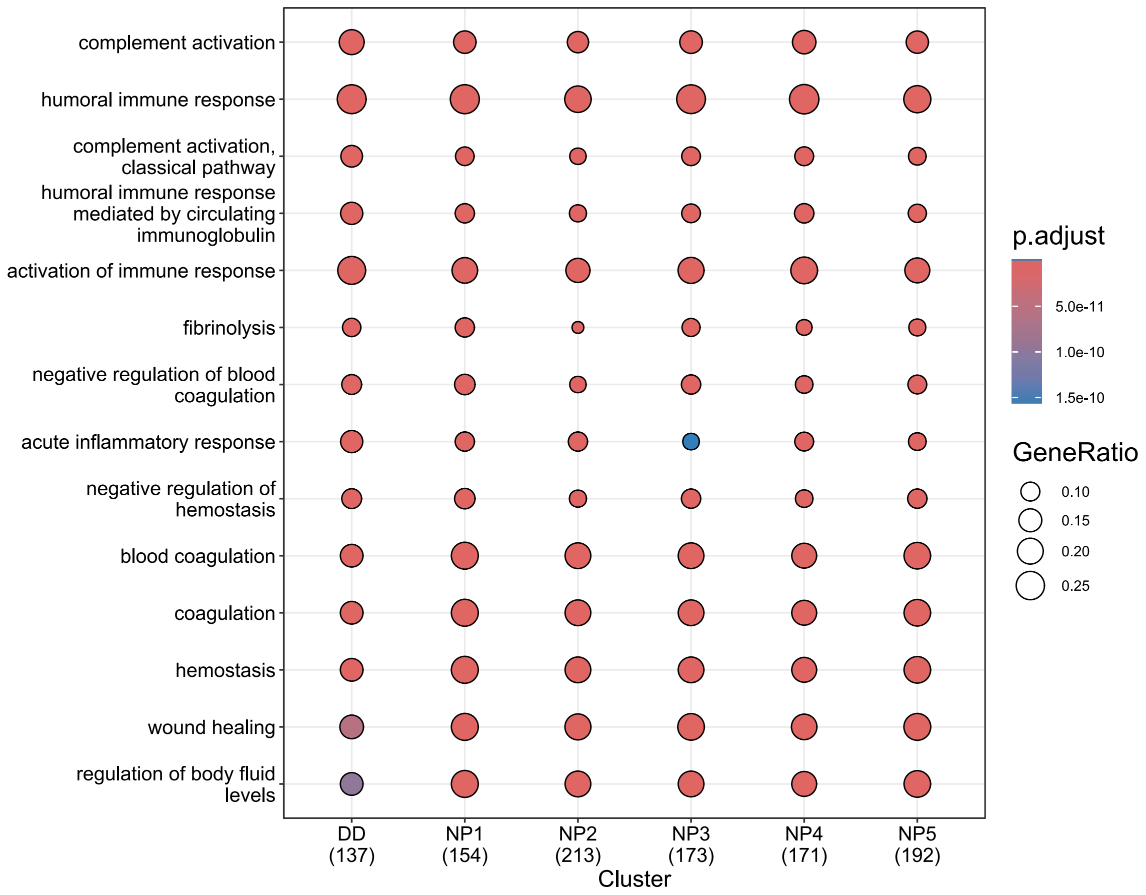


**Figure S16.** GO pathway enrichment analysis of different NPs and neat plasma (DD) by targeted DISPA. Fisher's exact test was applied and p-value <0.05 corrected by Benjamini–Hochberg.

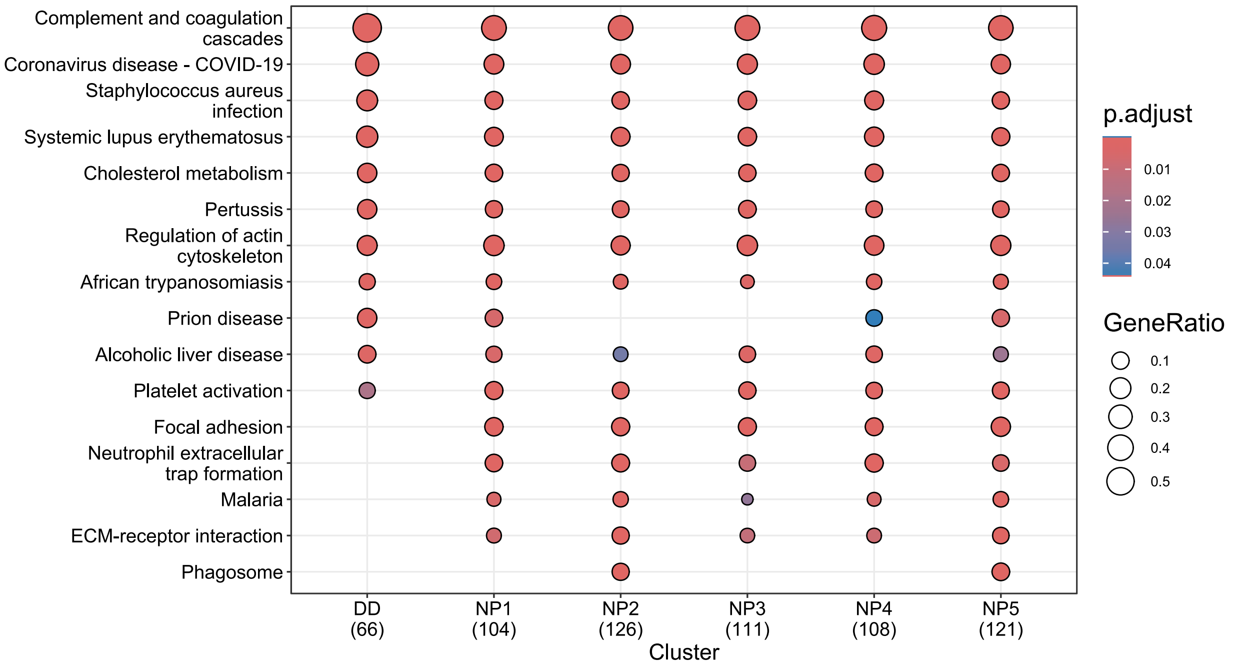


**Figure S17.** KEGG pathway enrichment analysis of different NPs and neat plasma (DD) by targeted DISPA. Fisher's exact test was applied and p-value <0.05 corrected by Benjamini–Hochberg.

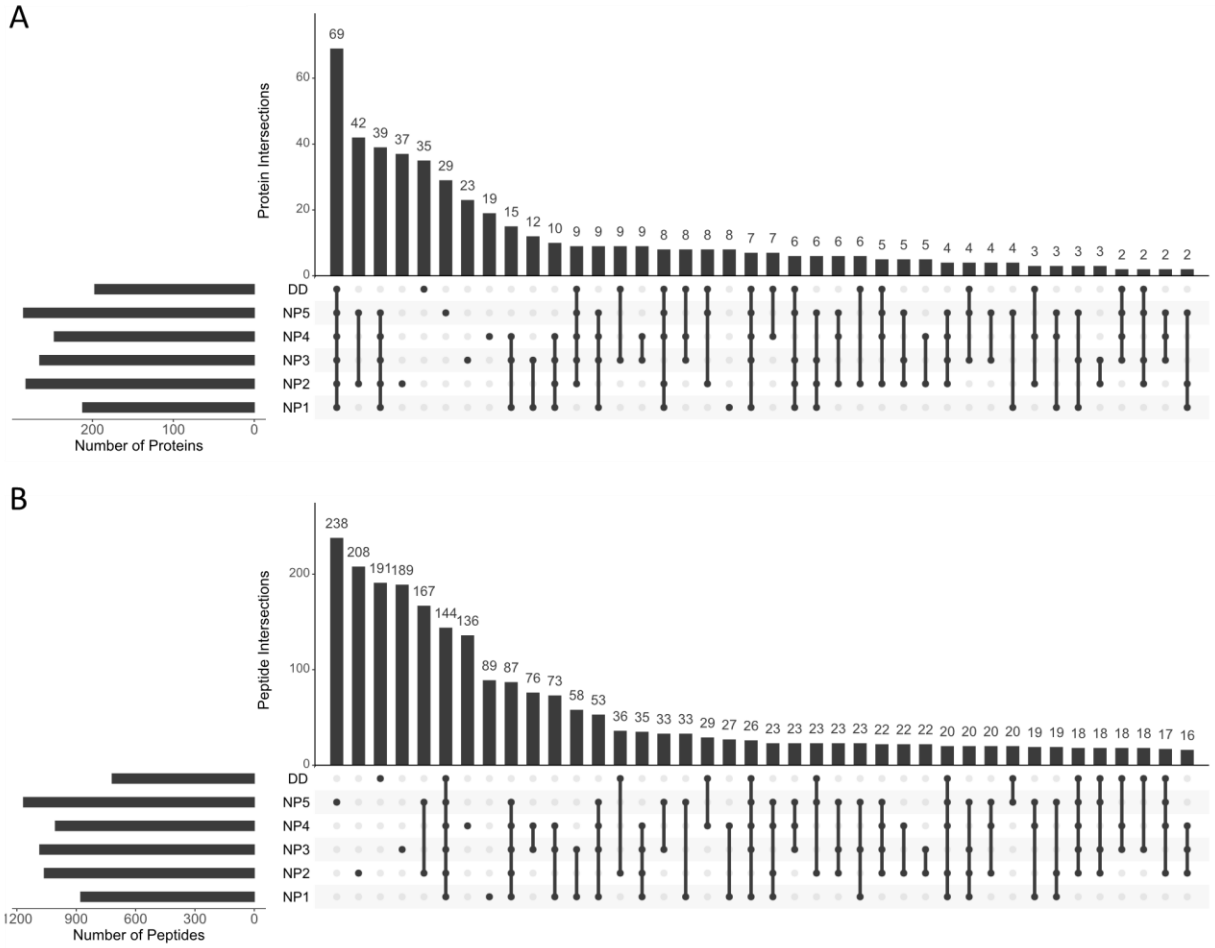


**Figure S18.** UpSet plot showing the overlap of identified proteins (A) and peptides (B) among 5 NPs and neat plasma by targeted DISPA. Protein groups were filtered for complete identifications across three replicates (n=3).

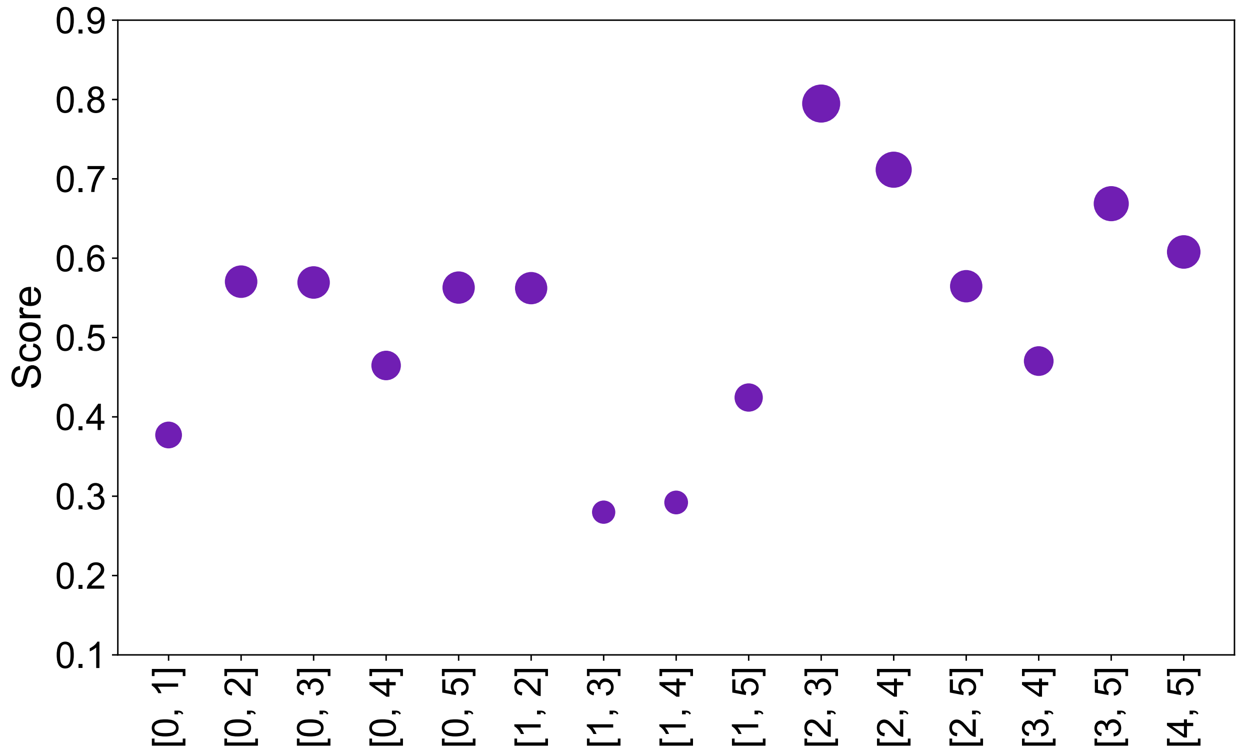


**Figure S19.** The score chart for each possible combination of the scenario for choosing two NPs. 0 represents neat plasma and 1 to 5 represents NP1 to NP5.

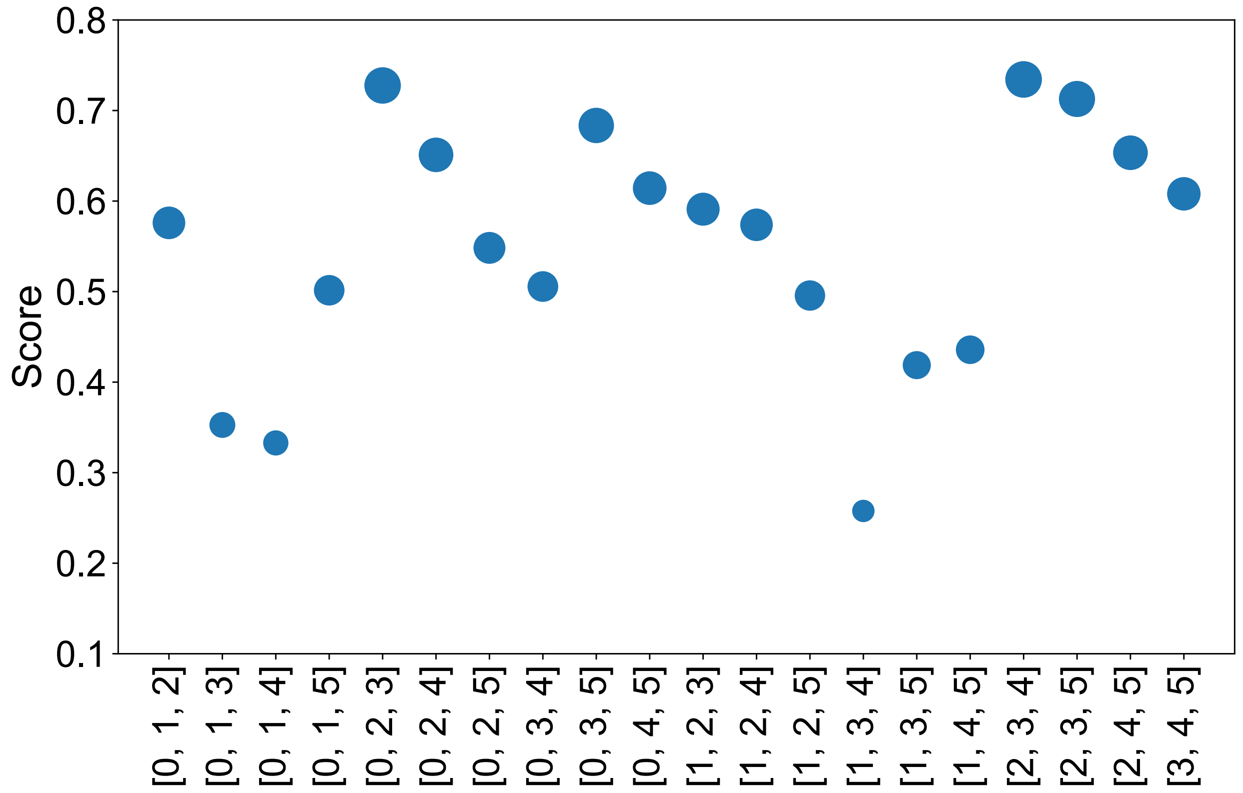


**Figure S20.** The score chart for each possible combination of the scenario for choosing three NPs. 0 represents neat plasma and 1 to 5 represents NP1 to NP5.

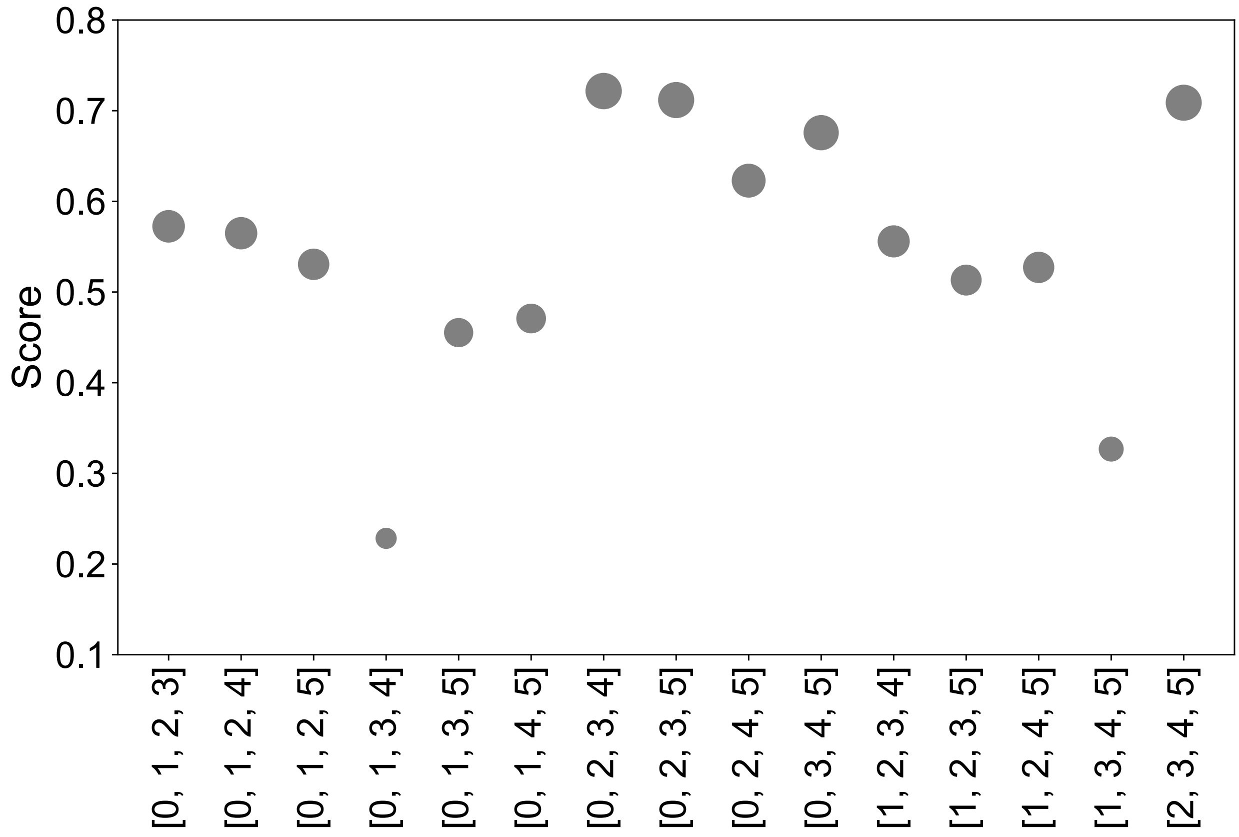


**Figure S21.** The score chart for each possible combination of the scenario for choosing four NPs. 0 represents neat plasma and 1 to 5 represents NP1 to NP5.
